# ClusterMine: a Knowledge-integrated Clustering Approach based on Expression Profiles of Gene Sets

**DOI:** 10.1101/255711

**Authors:** Hong-Dong Li, Yunpei Xu, Xiaoshu Zhu, Quan Liu, Gilbert S. Omenn, Jianxin Wang

## Abstract

**Motivation:** Clustering analysis is essential for understanding complex biological data. In widely used methods such as hierarchical clustering (HC) and consensus clustering (CC), expression profiles of all genes are often used to assess similarity between samples for clustering. These methods output sample clusters, but are not able to provide information about which gene sets (functions) contribute most to the clustering. So interpretability of their results is limited. We hypothesized that integrating prior knowledge of annotated biological processes would not only achieve satisfying clustering performance but also, more importantly, enable potential biological interpretation of clusters.

**Results:** Here we report ClusterMine, a novel approach that identifies clusters by assessing functional similarity between samples through integrating known annotated gene sets, *e.g.,* in Gene Ontology. In addition to outputting cluster membership of each sample as conventional approaches do, it outputs gene sets that are most likely to contribute to the clustering, a feature facilitating biological interpretation. Using three cancer datasets, two single cell RNA-sequencing based cell differentiation datasets, one cell cycle dataset and two datasets of cells of different tissue origins, we found that ClusterMine achieved similar or better clustering performance and that top-scored gene sets prioritized by ClusterMine are biologically relevant.

**Implementation and availability:** ClusterMine is implemented as an R package and is freely available at: www.genemine.org/clustermine.php

**Supplementary Information:** Supplementary data are available at Bioinformatics online.

## 1 Introduction

Clustering analysis for class discovery aims at identifying intrinsic groups of samples that display the same gene expression pattern. It is a very useful technique for understanding complex structures of biological data and is widely used in areas such as cancer subtyping and cell type classification. Commonly used general-purpose methods include hierarchical clustering (HC)(Mu, et al., 2017) and consensus clustering (CC) (Monti, et al., 2003; Wilkerson and Hayes, 2010). With the aid of these methods, one can computationally interrogate gene expression profiles and identify biologically meaningful clusters of samples that share the same pathological process. For example, based on graph theory, a modified version of the CC method was developed to explore clusters of genes in disease samples from CNS tumors, leukemia and lung cancers(Yu, et al., 2007). By clustering gene expression data, Bailey et al identified four subtypes of prostate cancers and identified opportunities for therapeutic development(Bailey, et al., 2016). Specifically, in the area of single cell RNA-sequencing (scRNA-seq) data analysis, several methods have been developed for cell type identification and discovery. These include single cell consensus clustering (SC3)(Kiselev, et al., 2017),SNN-cliq(Xu and Su, 2015), and SIMLR(Wang, et al., 2017).

At the core of clustering analysis approaches is the calculation of sample similarity. In the above-mentioned general-purpose or scRNA-seq specific approaches, the sample similarity is often calculated using holistic gene expression profiles or optionally a percentage (say 80%) of randomly selected genes, as in CC. Based on the common assumption that only a small percent of genes are differentially expressed between conditions such as disease or cell subtypes(Robinson and Oshlack, 2010), the resulting sample similarity may be noisy because its calculation involves a large number of genes that are not relevant to the biological differences under investigation. Therefore, the similarity based on holistic gene expression profiles may not accurately reflect the biological differences between sample clusters. Another limitation of the holistic gene expression-based similarity is that clustering results are often hard to interpret, because all genes are used and genes that contribute most to sample clustering are not prioritized and supplied to users.

To address these limitations, we propose a new general-purpose clustering method, called ClusterMine, for identifying clusters by integrating knowledge of gene sets that represent a biological function or pathway. The key feature of this method is that a local similarity matrix between samples is calculated for each biologically meaningful gene set, e.g. biological processes in Gene Ontology (GO), pathways in KEGG, and annotated gene sets in MSigDB(Subramanian, et al., 2005). The local similarity reflects the between-sample functional similarity with respect to the corresponding gene set. The cluster-separating ability of each gene set is also evaluated, therefore allowing prioritization of gene sets that are most likely to underlie clusters and enabling biological interpretability. Then, a global similarity matrix, denoted as **S**, is computed as the weighted sum of local similarity matrices resulting from individual gene sets. The global distance matrix can be calculated as **D**=**1**-**S**. Finally, clustering of samples is performed using the conventional hierarchical clustering method with **D** as the distance matrix. We tested ClusterMine on three cancer datasets, two scRNA-seq based cell differentiation datasets, one cell cycle dataset and two datasets of cells of different tissue origins, and compared its performance with the commonly used general-purpose Consensus Clustering approach(Monti, et al., 2003).

## 2 Materials and methods

### 2.1 Algorithm

Given a gene expression dataset of *n* samples and *p* genes, the goal of ClusterMine is not only to detect clusters of samples sharing the same gene expression profiles but also to prioritize functional gene sets that are most likely to underlie the separation between clusters. As depicted in **Figure 1**, two inputs are required to run ClusterMine, as detailed below:

1. Gene sets that represent known knowledge about biological functions of genes, which can be obtained from commonly used databases such as GO, KEGG, MSigDB(Subramanian, et al., 2005). Since both GO terms (biological processes, molecular function, cellular component) and KEGG pathways are included as part of the MSigDB database, we here only consider the gene sets annotated in MSigDB(Subramanian, et al., 2005). In this database, gene sets were organized into seven functionally different collections(Subramanian, et al., 2005): C1: positional gene sets; C2: curated gene sets; C3: motif gene sets; C4: computational gene sets; C5: GO gene sets; C6: oncogenic gene sets; C7: immunologic gene sets. To facilitate the use of these gene sets, we have downloaded them and built them into the ClusterMine R package. Users can choose which gene sets to analyze, without having to download them from the MSigDB website.
2. A gene expression matrix with *n* samples and *p* genes, which can be any gene expression data of interest. For example, they can be microarray or RNA-sequencing data, can be from bulk-sample or single cells, can be from healthy or disease samples.

**Fig. 1.**
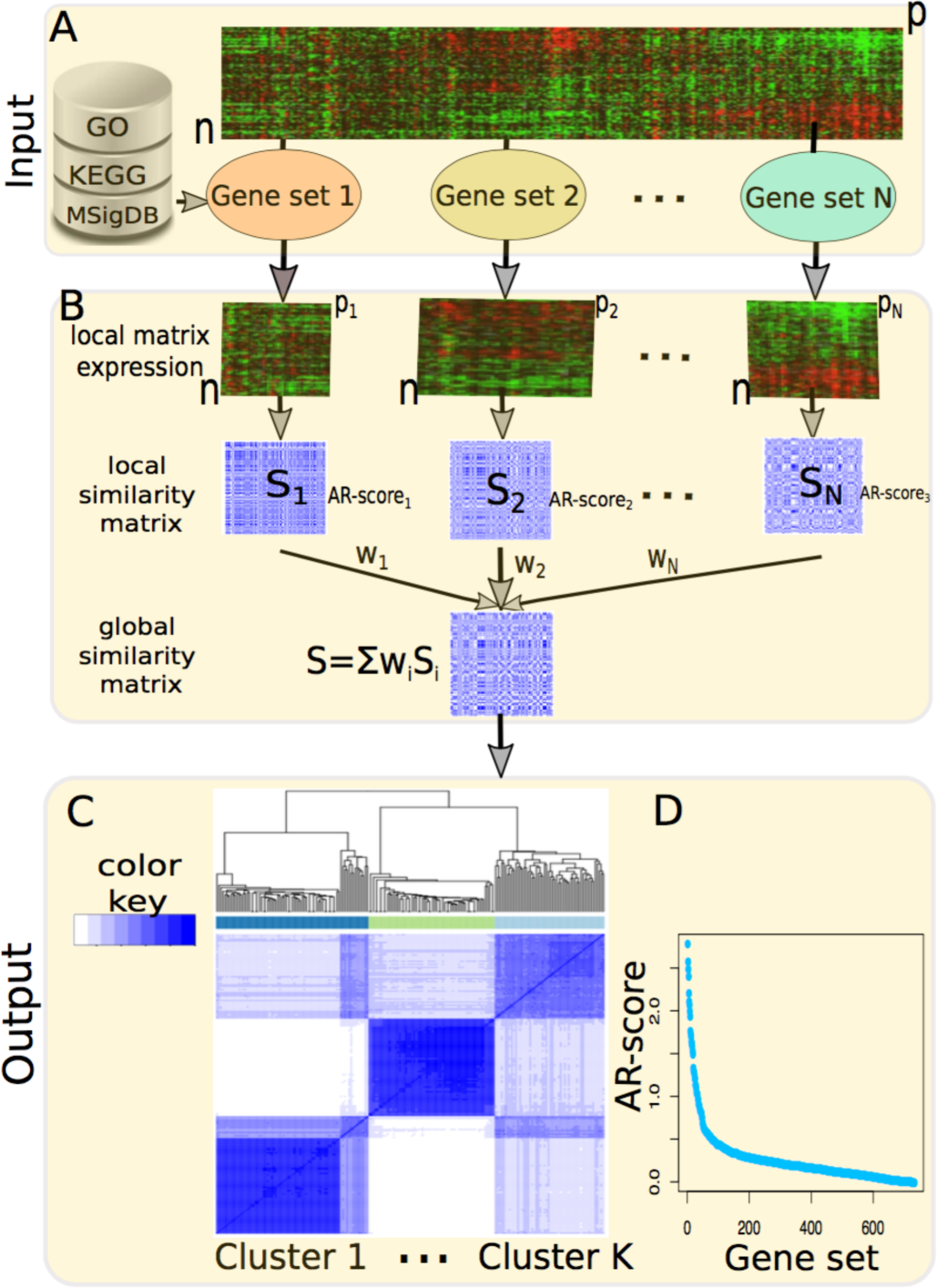
The schematic of ClusterMine. **A**. Two inputs of this method: (1) gene expression data and (2) known gene set database such as GO, KEGG and MSigDB. **B**. ClusterMine first calculates a local similarity matrix for each gene set, then computes a global similarity matrix, denoted as **S**, as the weighted sum of all the local similarity matrices. **C**. The global distance matrix **D,** calculated as **1-S,** is used for an agglomerative hierarchical clustering. **D**. An illustration of gene sets ranked by their average ratios (AR).

In addition to these two inputs, the number of clusters, denoted as K, needs be specified by the users based on their prior knowledge or optimization by computationally approaches to run ClusterMine. The algorithm of ClusterMine is detailed in the following.

First, for each gene set, we derive a local expression matrix, denoted as **X**_i_ of size *n*x*p*_i_, from the user-provided gene expression data by extracting only those genes in the gene set. *p*_i_ should be smaller than or equal to the number of all genes in the gene set, since some genes may not be expressed in the sample under investigation due to the known tissue-specific expression of genes. In single-cell studies, a large percentage of genes may be undetectable or undetected, leading to sparse datasets. Then, we perform hierarchical clustering on **X**_i_ using the *hclust* R function, and obtain the cluster membership of each sample using the *cutree* R function by setting the number of required clusters to user-specified K. Often, users test several K values and select an optimum based on expert knowledge. Based on the cluster membership, we can calculate a consensus matrix S_i_ of size *n* x *n* with its element at the *j*th row and *m*th column set to 1 if sample *j* and sample *m* belong to the same cluster and 0 otherwise.

Then, we evaluated the class separating performance of the gene set using Fisher discriminant analysis (FDA). For each of the K(K-1)/2 pairs of clusters, we calculated the ratio of between-group to within-group variance. We then use the average of all the K(K-1)/2 ratios as a score (AR-score) to measure the cluster-separating importance of gene sets. Since the local expression matrix **X**_i_ may have many more genes than samples, the co-variance matrix will be ill-conditioned and FDA models can not be accurately calculated. Also, over-fitting can easily occur when there are more genes than samples. To make FDA computable and reduce the risk of over-fitting, principal component analysis (PCA) is applied to reduce data dimension by decomposing **X**_i_ using PCA and choosing only the top ranked principal components that explain a predefined percentage, say 90%, of total variance as the input of FDA.

After analyzing all the N gene sets, we can obtain N similarity matrix **S**_i_, along with their AR-score_i_, i=1, 2, … N. Then, we calculate a weight for each gene set as the normalized AR-score_i_ using Formula (1):

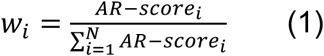

Further, a global similarity matrix is calculated as the weighted sum of the local similarity matrix as follows:

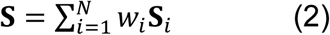

And the distance matrix between samples, denoted as **D**, is calculated as:

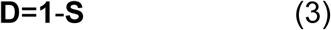

Then, conventional hierarchical clustering is performed with **D** as the distance matrix, and class membership of each sample is calculated according to the user-specified cluster number K. AR-scores are used to rank and prioritize gene sets that most likely underlie the separation between clusters.

### 2.2 Software implementation

ClusterMine was implemented as an R package. All the seven classes of gene sets in MSigDB (version 6.0)(Subramanian, et al., 2005), a rich compilation of gene sets about normal biology or cancers derived from GO, KEGG, literature mining, etc., were built into the package to facilitate users’ interrogation of gene expression data with respect to their functional gene sets of interest.

Three commonly used criteria to assess clustering performance, normalized mutual information (NMI), random index (RI) and adjusted random index (ARI), were also implemented and included in this package.

The software together with installation guidance and usage instructions is freely available at www.genemine.org/clustermine.php.

## 3 Results

We tested the performance of ClusterMine on three cancer datasets, two scRNA-seq based cell differentiation datasets, one cell cycle dataset and two datasets of cells of different tissue origins.

Gene sets in MSigDB were used in our study. Of the seven classes of gene sets (C1 to C7) in MSigDB(Subramanian, et al., 2005), we mainly considered three classes of gene sets (C2, C5 and C6) in our study. The reasons are that: (1) C2 represents curated gene sets from the biomedical literature, online databases such as KEGG and knowledge of domain experts, and are considered of high quality; (2) C5 collects gene sets in Gene Ontology which represent normal biology and are widely used in biological and bioinformatics studies; and (3) C6 is a collection of signatures of cellular pathways that are often dys-regulated in cancers and is therefore suitable to interrogate gene expression of cancer data. The reasons we do not consider C1, C3 and C7 are that: C1 (positional gene sets) and C3 (motif gene sets) do not describe biological functions, the focus of our study; C4 (computational gene sets) focuses on computationally derived gene sets and is thus less confident than experimentally validated or curated gene sets; C7 is specific to immunologic pathways. However, depending on interest and purpose, any gene set can be used for ClusterMine.

### 3.1 Cancer subtype identification

We tested the performance of ClusterMine in identifying subtypes in three cancer datasets (**Table 1**). (1) Garber dataset: this lung cancer dataset contains gene expression profiles of 39 adenocarcinoma, 13 squamous cell carcinoma and 5 normal control samples(Garber, et al., 2001). It was available at http://genome-www.stanford.edu/lung_cancer/adeno/data/lung_cluster.pcl. The downloaded data were already preprocessed; we calculated gene expression as the average of their probes. (2) Schlicker dataset: this colorectal cancer dataset contains two subtypes of adenocarcinoma samples (GDS4379). Subtype 1 has similar number of MSI and MSS samples, while Subtype 2 was enriched for MSS (Schlicker, et al., 2012). Gene expression values were summarized as the mean of probes; expression of each gene was median-centered. (3) Chung dataset: this breast cancer dataset(Chung, et al., 2017) consists of single-cell RNA sequencing data of 518 breast cancer and lymph node metastasis cells from 11 patients representing four subtypes of breast cancers: Luminal A (n=82), Luminal B (n=92), HER2 (n=161) and triple negative breast cancers (TNBC) (n=183). Since lowly expressed genes are noisy, genes with FPKM<1 in more than 80% samples were removed. Expression values were log_2_-transformed using *f*(x)=log_2_(x+1)(Eksi, et al., 2013); each gene value was median-centered.

**Table 1.** Clustering performance comparison of ClusterMine to Consensus Clustering (CC)(Wilkerson and Hayes, 2010) based on normalized mutual information (NMI) and random index (RI).

| Datasets | NMI |  |  |  | RI |  |  |  |
| --- | --- | --- | --- | --- | --- | --- | --- | --- |
|  | CC | ClusterMine |  |  | CC | ClusterMine |  |  |
|  | --- | C2 | C5 | C6 | --- | C2 | C5 | C6 |
| *Cancer datasets* |  |  |  |  |  |  |  |  |
| Garber | 0.433 | 0.488 | 0.427 | 0.515 | 0.650 | 0.662 | 0.618 | 0.692 |
| Schlicker | 0.429 | 0.429 | 0.460 | 0.400 | 0.725 | 0.725 | 0.748 | 0.703 |
| Chung | 0.295 | 0.328 | 0.304 | 0.355 | 0.596 | 0.647 | 0.693 | 0.669 |
| *Cell differentiation* |  |  |  |  |  |  |  |  |
| Yeo | 0.846 | 0.975 | 0.975 | 0.975 | 0.923 | 0.993 | 0.993 | 0.993 |
| Kolod | 1.000 | 0.992 | 1.000 | 1.000 | 1.000 | 0.998 | 1.000 | 1.000 |
| *Cell cycle* |  |  |  |  |  |  |  |  |
| Sasagawa | 0.307 | 0.511 | 0.652 | 0.281 | 0.498 | 0.684 | 0.810 | 0.636 |
| *Tissue mixture* |  |  |  |  |  |  |  |  |
| Pollen | 0.907 | 0.941 | 0.942 | 0.937 | 0.958 | 0.979 | 0.980 | 0.979 |
| Ramskold | 0.939 | 0.939 | 0.928 | 0.939 | 0.968 | 0.968 | 0.960 | 0.968 |

For the Garber data (lung cancer), we first compared the performance of our method with state-of-the-art Consensus Clustering (CC) in terms of NMI and RI. The number of clusters for both methods was set to 3, corresponding to the three subclasses of sample. The NMI and RI by CC and ClusterMine (based on C2, C5 and C6) are listed in **Table 1**. Compared to CC, ClusterMine achieved a 5%-8% positive increment of NMI on C2 and C6 datasets, but decreased by 0.6% on C5, with similar observations using the RI criterion. Thus, ClusterMine achieved on average a better performance than CC.

For ClusterMine, the highest NMI or RI was achieved on the C6 class of gene sets, suggesting that, within this dataset, the subtyping of lung cancers could be better explained by known cancer-related dys-regulated pathways. Note that all the NMI values are low, which might imply the existence of gene sets that contribute to the heterogeneity of lung cancers but remain to be identified. The clustering heatmap based on C6 is shown in the left of **Figure 2A**, with the weights of gene sets that are not only in C6 but also expressed displayed in the right of **Figure 2A**. For example, the highest-scored gene set is EGFR_UP.V1_UP (AR-score=2.93), which contains a total of 193 genes. Mutation or amplification of EGFR is known to cause lung cancers and other cancers(Bethune, et al., 2010). Further, we performed GO enrichment analysis using GoTermFinder(Boyle, et al., 2004), and found that this set of genes is enriched in cancer-related biological processes including apoptotic process (P-corrected=1.16×10^-9^) and programmed cell death (P-corrected=1.48×10^-9^). E2F1_UP.V1_DN (AR-score=2.67) and RAF_UP.V1_UP (AR-score=2.11) are the top second and third gene sets, in which the genes *E2F1* and *RAF* also have been reported as associated with lung cancers(Hung, et al., 2012; Yu, et al., 2002). These results indicate that gene sets prioritized by ClusterMine are biological meaningful.

**Fig. 2.**
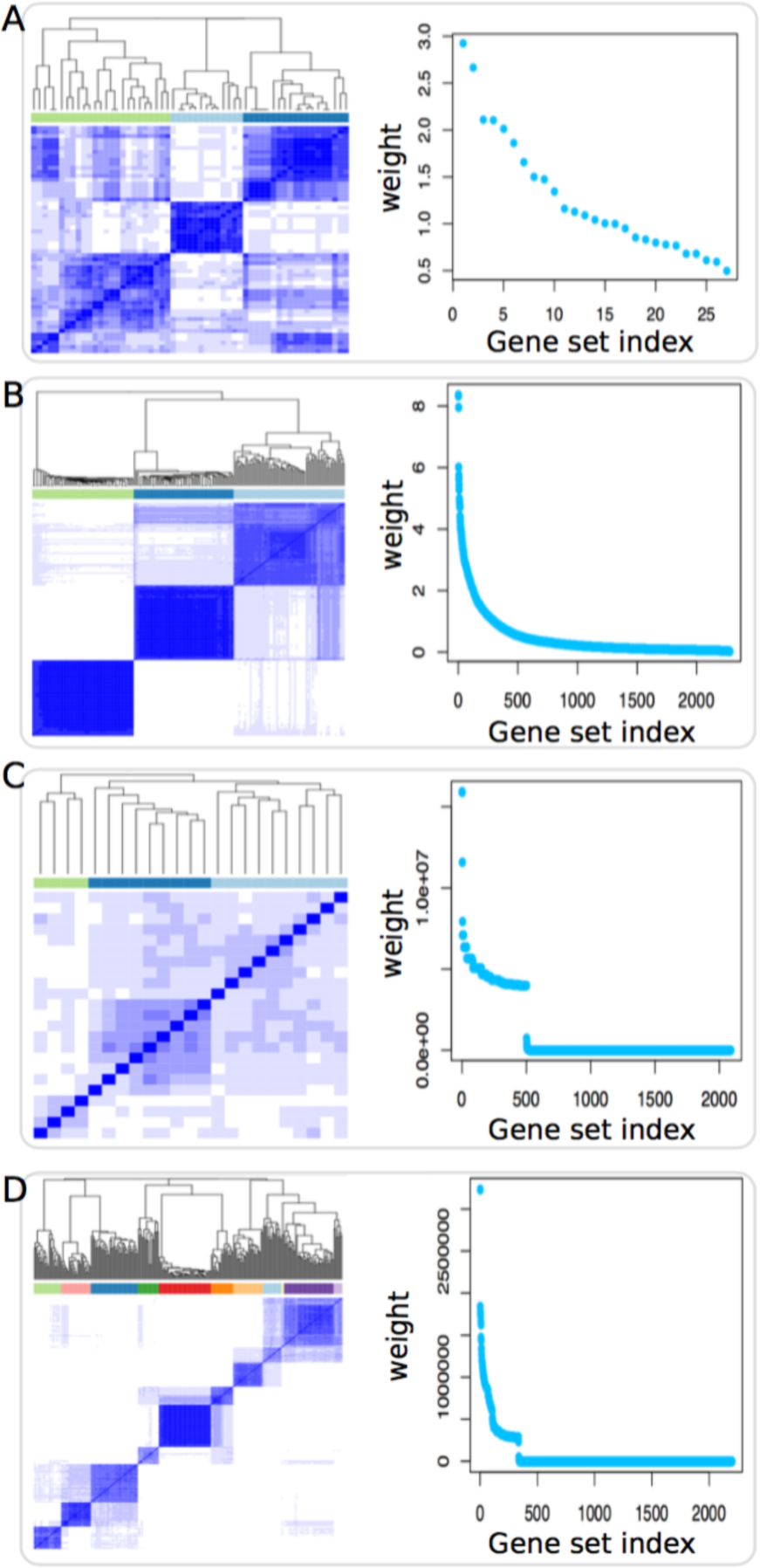
Example clustering heatmaps and the corresponding gene set weight plot based on the gene set associated the highest NMI. One example from each dataset category (Table 1) was shown. (**A**) the Garber dataset based on C6 gene set; (**B**) the Yeo dataset based on C2 gene set; (**C**) the Sasagawa dataset based on C5 gene sets; (**D**) the Pollen dataset based on C5 gene sets. The weight is a measurement of the cluster-separating ability of gene sets. The higher the weight is, the more likely the corresponding gene set underlies the clustering of samples.

For the Schlicker dataset (colorectal cancer), the best performance was achieved by our method on the C5 gene sets, from which both NMI or RI values are 2%-3% higher compared to CC (**Table 1**). The performance on C2 keeps the same as that of CC and the performance on C6 showed a slight decrease. These results indicate ClusterMine is a promising method for clustering with overall better performance over CC. In addition to the satisfying clustering performance, we also found that the top-scored gene sets by our method are disease-relevant. For example, the highest-scored gene set in C5 is GO_STRUCTURAL_CONSTITUENT_OF_RIBOSOME (AR-score = 2.18). Our literature survey showed that ribosome proteins, which are key constituents of ribosomes, were associated with colorectal cancers(Goudarzi and Lindstrom, 2016; Lai and Xu, 2007). Another example is the second-ranked gene set GO_RESPONSE_TO_INSULIN (AR-score = 1.74). Insulin, directly involved in this gene set, was reported to be involved in the development of colon cancer(Giovannucci, 2001), implying the biological relevance of this gene set.

Overall, for the Schlicker dataset, we found that ClusterMine does not show clear advantage over CC in prediction performance. Such results may be an indication that annotated gene sets cannot fully represent the dys-regulated pathways between different conditions. In this case, CC would be a better choice over ClusterMine.

For the Chung dataset (breast cancer), we found that our method achieves better clustering performance than CC, in terms of higher NMI and RI values (**Table 1**). Based on NMI values, it can be seen that ClusterMine achieves the best performance on C6. The top ranked gene sets are TBK1.DF_DN (AR-score=1.52) and TBK1.DF_UP (AR-score=0.45), corresponding to the protein-coding gene *TBK1* (TANK Binding Kinase 1). *TBK1* has been associated with breast cancer, and knockdown of *TBK1* suppressed growth of human HER2+ breast cancer cells(Jiang, et al., 2014), supporting the relevance of this gene set to breast cancers.

Further, for each of the above three datasets, we sorted all gene sets in C2, C5 and C6 by their AR-scores, and provided the top 5 ranked gene sets in **Table 2**. These gene sets are most likely associated with the disease under investigation, and may be a useful resource for the community.

**Table 2.**
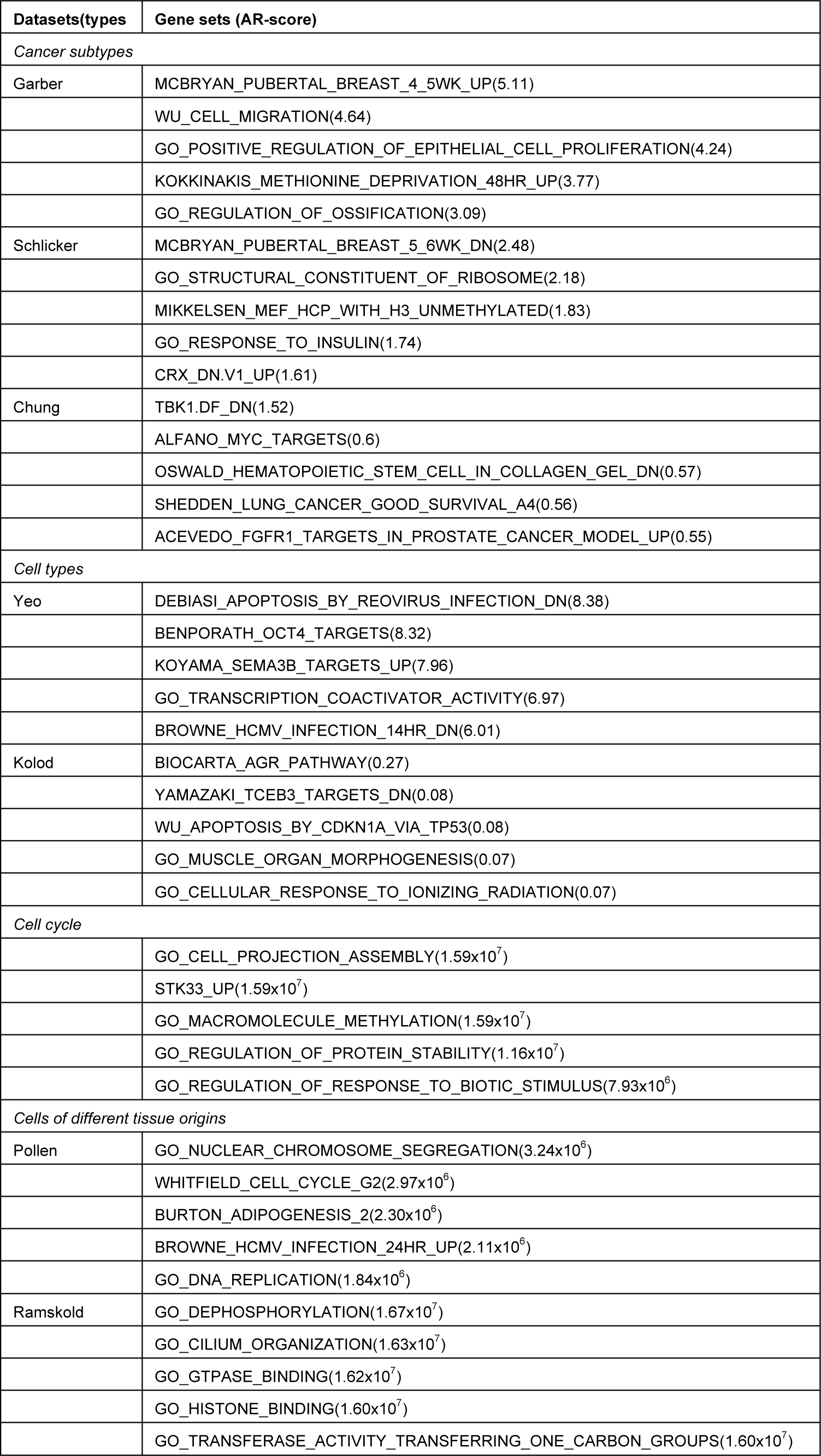
The top 5 scored gene sets of each dataset along with their AR-scores shown in parentheses.

| Datasets(types) | Gene sets (AR-score) |
| --- | --- |
| *Cancer subtypes* |  |
| Garber | MCBRYAN_PUBERTAL_BREAST_4_5WK_UP(5.11) |
|  | WU_CELL_MIGRATION(4.64) |
|  | GO_POSITIVE_REGULATION_OF_EPITHELIAL_CELL_PROLIFERATION(4.24) |
|  | KOKKINAKIS_METHIONINE_DEPRIVATION_48HR_UP(3.77) |
|  | GO_REGULATION_OF_OSSIFICATION(3.09) |
| Schlicker | MCBRYAN_PUBERTAL_BREAST_5_6WK_DN(2.48) |
|  | GO_STRUCTURAL_CONSTITUENT_OF_RIBOSOME(2.18) |
|  | MIKKELSEN_MEF_HCP_WITH_H3_UNMETHYLATED(1.83) |
|  | GO_RESPONSE_TO_INSULIN(1.74) |
|  | CRX_DN.V1_UP(1.61) |
| Chung | TBK1.DF_DN(1.52) |
|  | ALFANO_MYC_TARGETS(0.6) |
|  | OSWALD_HEMATOPOIETIC_STEM_CELL_IN_COLLAGEN_GEL_DN(0.57) |
|  | SHEDDEN_LUNG_CANCER_GOOD_SURVIVAL_A4(0.56) |
|  | ACEVEDO_FGFR1_TARGETS_IN_PROSTATE_CANCER_MODEL_UP(0.55) |
| *Cell types* |  |
| Yeo | DEBIASI_APOPTOSIS_BY_REOVIRUS_INFECTION_DN(8.38) |
|  | BENPORATH_OCT4_TARGETS(8.32) |
|  | KOYAMA_SEMA3B_TARGETS_UP(7.96) |
|  | GO_TRANSCRIPTION_COACTIVATOR_ACTIVITY(6.97) |
|  | BROWNE_HCMV_INFECTION_14HR_DN(6.01) |
| Kolod | BIOCARTA_AGR_PATHWAY(0.27) |
|  | YAMAZAKI_TCEB3_TARGETS_DN(0.08) |
|  | WU_APOPTOSIS_BY_CDKN1A_VIA_TP53(0.08) |
|  | GO_MUSCLE_ORGAN_MORPHOGENESIS(0.07) |
|  | GO_CELLULAR_RESPONSE_TO_IONIZING_RADIATION(0.07) |
| *Cell cycle* |  |
| | GO_CELL_PROJECTION_ASSEMBLY($1.59 \times 10^7$) |
| | STK33_UP($1.59 \times 10^7$) |
| | GO_MACROMOLECULE_METHYLATION($1.59 \times 10^7$) |
| | GO_REGULATION_OF_PROTEIN_STABILITY($1.16 \times 10^7$) |
| | GO_REGULATION_OF_RESPONSE_TO_BIOTIC_STIMULUS($7.93 \times 10^6$) |
| *Cells of different tissue origins* |  |
| Pollen | GO_NUCLEAR_CHROMOSOME_SEGREGATION($3.24 \times 10^6$) |
| | WHITFIELD_CELL_CYCLE_G2($2.97 \times 10^6$) |
| | BURTON_ADIPOGENESIS_2($2.30 \times 10^6$) |
| | BROWNE_HCMV_INFECTION_24HR_UP($2.11 \times 10^6$) |
| | GO_DNA_REPLICATION($1.84 \times 10^6$) |
| Ramskold | GO_DEPHOSPHORYLATION($1.67 \times 10^7$) |
| | GO_CILIUM_ORGANIZATION($1.63 \times 10^7$) |
| | GO_GTPASE_BINDING($1.62 \times 10^7$) |
| | GO_HISTONE_BINDING($1.60 \times 10^7$) |
| | GO_TRANSFERASE_ACTIVITY_TRANSFERRING_ONE_CARBON_GROUPS($1.60 \times 10^7$) |

### 3.2 Cell type identification

We further applied ClusterMine for cell type identification in two scRNA-seq datasets. (1) Yeo dataset(Song, et al., 2017): single cell RNA-seq expression from 62 induced pluripotent stem cells (iPSC), 69 neural progenitor cells (NPCs), and 60 motor neurons (MNs). (2) Kolod dataset: scRNA-seq data from 704 pluripotent mesenchymal stem cells under three different culture conditions(Kolodziejczyk, et al., 2015). Gene expression was quantified as RPKM for these two datasets. Since lowly expressed genes are noisy, genes with RPKM or TPM<1 in more than 80% samples were removed. Expression values were log_2_-transformed using *f*(x)=log_2_(x+1)(Eksi, et al., 2013). Each gene was median-centered.

For the Yeo dataset, we found that ClusterMine achieves much higher accuracies in the identification of different cell types, with NMI and RI improved by 12.9% and 7%, respectively over the Consensus Clustering method (**Table 1**). As an illustration, we provided the heatmap of the clustering of all the cells based on the curated gene sets of C2 in **Figure 2B** (left), with the AR-scores of the gene sets shown alongside in **Figure 2B** (right). Of interest, we found that top-scored gene sets are related to stem cell differentiation. For example, the first-ranked is DEBIASI_APOPTOSIS_BY_REOVIRUS_INFECTION_DN (AR-score=8.38). GO enrichment shows that this gene set is mainly enriched in functions about development and cell differentiation, such as development process (P-corrected=1.43×10^-9^), anatomic structure development (P-corrected=1.86×10^-8^) and cell differentiation (P-corrected=8.82×10^-7^). The second ranked gene set is BENPORATH_OCT4_TARGETS (AR-score=8.32), which is enriched in tissue development (P-corrected=1.13×10^-14^), cellular developmental process (P-corrected=2.57×10^-12^), cell differentiation (P-corrected=2.39×10^-12^) etc. These results provide evidence that ClusterMine is a promising tool for clustering analysis, achieving high accuracies and enabling biological interpretation.

For Kolod dataset, the NMI and RI values from both methods are very close or equal to 1, suggesting that gene expression profiles of the cells under different conditions show large differences and can be easily distinguished from each other. Top gene sets were also found to be biologically relevant. Taking as an example the first-ranked gene set in C2 (BIOCARTA_AGR_PATHWAY with AR-score=0.27), it is enriched for GO biological process such as signal transduction by protein phosphorylation (P-corrected=3.58×10^-7^), which is known to play key roles in cell differentiation(Bononi, et al., 2011).

Taken together, the above results support that ClusterMine is a promising approach for clustering analysis to identify cell types with high accuracies and provide information about gene sets are likely to be relevant to cell type differentiation.

### 3.3 Cell cycle identification

We obtained a cell cycle dataset from Sasagawa(Sasagawa, et al., 2013). This dataset contains single-cell Quartz-seq data of mouse ES cells in different cell-cycle phases. A total of 23 cells were analyzed in three phases (G1: n=8, S: n=7, G2/M: n=8). Gene expression was quantified as FPKM in their work(Sasagawa, et al., 2013). In our analysis, genes with FPKM<1 in more than 80% samples were removed. Genes were log_2_-transformed and median centered.

The clustering performance of CC and ClusterMine is shown in **Table 1**. For NMI, our method outperformed CC on the C2 and C5 classes of gene sets, with slightly lower NMI values on C6. In terms of RI values, our method showed significantly better results on all the three gene sets. This result suggests that our method is promising in distinguishing cells at different phases.

The top ranked gene set is GO_CELL_PROJECTION_ASSEMBLY (AR-score=1.59 x10^7^), a gene set containing 264 genes. Enrichment analysis for this gene set using GoTermFinder(Boyle, et al., 2004) showed it was significantly enriched in a large number of biological processes, including, but not limited to, cellular component assembly (P-corrected = 8.98×10^-145^) and microtubule-based process (3.86×10^-49^), which are known to be involved in cell cycle(Forth and Kapoor, 2017; Hernandez-Verdun, 2011). The second ranked is GO_MACROMOLECULE_METHYLATION (AR-score=1.59 x10^7^; note that this score is actually lower than the first ranked one but the difference can not be reflected with two decimals), which was shown to be related to cell cycle(Barwick, et al., 2016; Vandiver, et al., 2015). These results indicate that our method was able to prioritize gene sets that are relevant to the biologic problem of interest. As a resource, we sorted all gene sets in C2, C5 and C6, and provided the top five gene sets in **Table 2**.

To intuitively visualize the clustering of the samples, the heatmap of cluster together with the gene set weight plot was displayed in **Figure 2C**.

### 3.4 Cell mixtures of different tissue origins

Two datasets containing cells with different tissue origins were used. (1) Pollen dataset: gene expression profiles of 11 cell populations, including skin cells, pluripotent stem cells, blood cells, and neural cells(Pollen, et al., 2014). (2) Ramskold dataset: generated using SMART-seq, a variant single cell RNA-seq protocol (Ramsköld, et al., 2012), this dataset contains gene expression profiles of cells from different tissues: human embryonic stem cells (n=8), putative melanoma circulating tumor cells (n=6), melanoma cell lines SKMEL5 (n=4), prostate cancer cell lines LNCap (n=4), bladder cancer cell line T24 (n=4), PC3 (n=4), UACC257 (N=3). For both datasets, genes with FPKM<1 in more than 80% samples were removed. Genes were log2-transformed and median centered. K was set to 11, corresponding to the 11 cell populations.

For the Pollen dataset, the NMI by the CC method is 0.907, which is consistently lower than that from ClusterMine based on C2 (0.941), C5 (0.942) and C6 (0.937) (**Table 1**). The RI values of ClusterMine are also higher than that of CC. Similarly, we found that functions of the top ranked gene sets are biologically relevant. Examples include gene sets in C2 such as WHITFIELD_CELL_CYCLE_G2 (AR-score=2.9×10^6^) and BURTON_ADIPOGENESIS_2 (AR-score=2.3×10^6^), which were reported to be associated with cell fate and cell differentiation, respectively(Boward, et al., 2016). The clustering heatmap for this dataset was provided in **Figure 2D**, giving an intuitive view of how these samples clustered.

For the Ramskold dataset, ClusterMine based on C2 and C6 achieved the same performance as CC in terms of both NMI and RI, while the performance on C5 showed a slight (1%) decrease. Overall, the results from both methods are comparable. For these data, we found that prioritized gene sets were of biological relevance. Taking gene sets in C2 as an example, the top ranked is FORTSCHEGGER_PHF8_TARGETS_UP (AR-score=1.56×10^7^. Using GoTermFinder(Boyle, et al., 2004), this gene set was enriched in many biological processes, such as animal organ morphogenesis (P-corrected = 7.54×10^-5^), tissue development (P-corrected = 5.0×10^-4^), cell differentiation (P-corrected = 3.0×10^-3^), which are likely to be associated with cell differentiation into different types. The second ranked is BROWNE_HCMV_INFECTION_16HR_UP (AR-score=1.56×10^7^; note that this score is actually lower than the first ranked one but the difference can not be reflected with two decimals), which is also enriched in biological processes such as anatomical structure development (1.13×10^-9^) and developmental process (1.22×10^-9^). These results again suggest that gene sets prioritized by our method are biologically meaningful. Also, we sorted all gene sets in C2, C5 and C6, and provided the top five gene sets in **Table 2**.

## 3 Discussion

Clustering samples based on their gene expression profiles plays a key role in understanding complex biological data by grouping samples into clusters that show more nearly homogeneous biological pathways. Existing approaches like ConsensusClustering often use the full gene expression profiles (or a randomly sampled proportion, say 80%, of them) for clustering analysis and do not report which gene sets most likely underlie the differences between clusters(Wilkerson and Hayes, 2010), thus leading to limited interpretability of the resulting clustering.

We developed ClusterMine as a new tool to perform cluster discovery, which features in the integration of known gene sets representing our knowledge about gene functions. Another feature of this method is its ability to single out gene sets that contribute most to the separation between clusters by using Fisher discriminant analysis. This method is implemented as an R package. All the gene sets in MSigDB(v6.0) were built into this package to facilitate use of our method.

By analyzing two cancer datasets and three scRNA-seq datasets, we found that ClusterMine showed similar or better performance in terms of NMI (normalized mutual information) and RI (random index) compared to the commonly used Consensus Clustering (CC) approach. In addition to providing cancer subtypes or cell types in the given dataset, our method also analyzes gene sets that are biologically relevant. This feature is essential because it can guide us to infer the biology behind clustering and to propose hypotheses for further testing.

As can be seen from our analysis, different gene sets could lead to different results. Therefore, the performance of ClusterMine is dependent on the selection of gene sets. In general, users can choose to test gene sets of their interest. In the case of no prior knowledge, it is our recommendation that C2 (curated gene sets) and C5 (commonly used gene sets in Gene Ontology) be tested. In the case of cancer-related datasets, C6 (disturbed pathways in cancers) can be tested. In special cases of studying immunity-related biological questions, C7 can be used. As an option, users can also combine all the seven gene sets as input for ClusterMine, but using all gene sets does not necessarily or reliably improve clustering performance, since many gene sets irrelevant to the clustering are involved in the assessment of between-sample similarity and thus introduce noise. Here, feature selection approaches, though beyond the scope of this report, can be used to select a subset from all the combined gene sets, followed by feeding the selected subsets again to ClusterMine for clustering analysis.

ClusterMine also has limitations. Firstly, since gene products, especially in higher organisms such as humans, are a mixture of isoforms generated by the alternative splicing mechanism(Li, et al., 2014; Tapial, et al., 2017; Xiong, et al., 2015), isoform-level gene expression would promise more accurate clustering. Because our method is dependent on function annotations and current functions are mainly only available at the gene level, ClusterMine can only analyze gene-level expression data and is not able to directly take isoform-level expression as input. Other methods that are not dependent on gene function annotation such as the Consensus Clustering (CC) method can analyze both gene‐ and isoform-level expression data. With the accumulation of experimentally validated isoform function annotations or using predicted isoform functions(Li, et al., 2015; Li, et al., 2015), ClusterMine can then be extended to isoform-level data analysis. Secondly, compared to methods such as CC, ClusterMine is computationally more extensive since it computes a similarity matrix for each gene set. Taking the C2 collection of 4378 gene sets as example, a total of 4378 similarity matrices are needed to be computed. As a result, ClusterMine often runs slower, though the time to run ClusterMine is not long (seconds to minutes in our analyzed data).

Summing up, ClusterMine is a competitive and complementary tool for clustering analysis. We expect that it will find many more applications in bioinformatics and biomedical research.

